# RGBM: Regularized Gradient Boosting Machines for the Identification of Transcriptional Regulators of Discrete Glioma Subtypes

**DOI:** 10.1101/132670

**Authors:** Raghvendra Mall, Luigi Cerulo, Khalid Kunji, Halima Bensmail, Thais S. Sabedot, Houtan Noushmehr, Antonio Iavarone, Michele Ceccarelli

## Abstract

The transcription factors (TF) which regulate gene expressions are key determinants of cellular phenotypes. Reconstructing large-scale genome-wide networks which capture the influence of TFs on target genes are essential for understanding and accurate modelling of living cells. We propose RGBM: a gene regulatory network (GRN) inference algorithm, which can handle data from heterogeneous information sources including dynamic time-series, gene knockout, gene knockdown, DNA microarrays and RNA-Seq expression profiles. RGBM allows to use an a priori mechanistic of active biding network consisting of TFs and corresponding target genes. RGBM is evaluated on the DREAM challenge datasets where it surpasses the winners of the competitions and other established methods for two evaluation metrics by about 10-15%.

We use RGBM to identify the main regulators of the molecular subtypes of brain tumors. Our analysis reveals the identity and corresponding biological activities of the master regulators driving transformation of the G-CIMP-high into the G-CIMP-low subtype of glioma and PA-like into LGm6-GBM, thus, providing a clue to the yet undetermined nature of the transcriptional events driving the evolution among these novel glioma subtypes.

RGBM is available for download on CRAN at https://cran.rproject.org/web/packages/RGBM/index.html

## 1 Introduction

Changes in environment and external stimuli lead to variations in gene expression which accordingly adapts for proper functioning of living systems. However, abnormalities in this tightly co-ordinated process are precursors to many pathologies. A vital role is played by the transcription factors (TF), which are proteins that bind to the DNA in the regulatory regions of specific target genes. These TFs can then repress or induce the expression of target genes. Many such transcriptional regulations have been discovered through traditional molecular biology experiments and several of these high-quality mechanistic regulatory interactions have been documented into TF-target gene databases [1].

With the availability of high-throughput experimental techniques for efficiently measuring gene expression, such as DNA micro-arrays and RNA-Seq, the aim now is to design computational models for gene regulatory network (GRN) [2] inference at genomic scale. The accurate reconstruction of GRNs from diverse gene expression information sources is one of the most important problems in biomedical research [3]. This is primarily because precisely reverse-engineered GRNs can reveal mechanistic hypotheses about differences between phenotypes and sources of diseases [1], which can ultimately help in the identification of therapeutic targets. The problem of inferring GRN from heterogeneous information sources such as dynamic time-series data, gene knockout, gene knockdown, protein-protein interactions etc. is one of the most actively pursued problem in computational biology [4] and has resulted in several DREAM challenges.

This problem is complicated by the noisy and high-dimensional nature of the data [5] which obscures the regulatory network with indirect connections. Another common challenge is to identify and model the non-linear relationships among the genes in the presence of relatively few samples compared with the total number of genes (i.e. *n* ≪ *p*). A majority of the computational techniques model the expression of an individual target gene as either a linear or nonlinear function of the expression levels of TFs. These modelling techniques use inductive logic to build a regulatory network which explains the interactions between the TFs and the target genes. Hence, these methods do not explain mechanistically how TFs interact with regulated genes, rather identify strong candidate interactions using the expression data. In this paper, we follow this traditional principle and identify candidate transcriptional regulations inferred from the gene expression data.

Several statistical methods have been proposed for inferring GRNs including TwixTwir [6] that uses a double two-way t-test to score transcriptional regulations, null-mutant z-score algorithm [7] which is based on z-score transformed knockout expression data, least angle regression based method TIGRESS [8] captures linear interactions, bayesian networks [9, 10, 11, 12] which employ probabilistic graphical models that analyze multivariate joint probability distributions over each target gene and a formal method based approach [13]. Several information theoretic methods have also been developed for reverse-engineering GRNs such as [14] and CLR [15]. The mutual information based technique ARACNE [16, 17] works on reducing the indirect edges by means of techniques such as Data Processing Inequality.

Of-late several machine-learning techniques have been utilized for construction of GRNs. Here the network inference problem is formulated as a series of supervised learning tasks, where each target gene is considered as a dependent variable and the list of TFs are the corresponding independent variables. The goal of the machine learning model is to provide a ranked list of TFs for each target gene as it captures the relative importance of the transcriptional regulations. Hence, several non-linear machine learning techniques including gradient boosting machines (GBM) based methods like ENNET [18] and OKVAR-Boost [19], random-forest based methods such as iRafNet [20] and GENIE3 [21] have been successfully employed for the task of GRN inference. These methods score important TFs, utilizing the embedded relative importance measure of input independent variables as a ranking criterion.

Many of these methods have participated in DREAM challenges for inferring GRN and achieved high performance w.r.t. area under precision-recall (*AU*_*pr*_) curve and area under the receiver operating characteristic (*AU*_*roc*_) curve. A characteristic of real world gene regulatory networks is that there are very few transcription factors (TF) that regulate each target [4] and there are a few targets which are not regulated by any TF [4]. One drawback of these machine learning techniques [18, 19, 20, 21] is that due to lack of regularization a large number of TFs gain variable importance and have connections with an individual target gene. This results in several false positive connections in the final inferred GRN. Moreover, these methods cannot identify genes with no incoming edges i.e. genes that are not regulated by any TF or are upstream regulators.

In this paper, we propose a method whose core model is boosting of regression stumps. The proposed method allows the user to specify *a priori* mechanistic active binding network (ABN) of TFs and target genes. In the presence of an ABN, the reconstructed network is a subgraph of it.

The reverse-engineering procedure infers an initial set of transcriptional regulations from the expression data (from heterogeneous information sources) using boosting of regression stumps (GBM). The obtained (TF-target) edge weights are then refined by any additional data such as knockout information. This refinement procedure is similar to that used in null-mutant z-score method [7] and ENNET algorithm [18].

In order to address the problems suffered by tree-based modeling techniques, we employ a notion used for identifying the corner of the L-curve criterion [22] in Tikonov regularization [23] on the edge weights (variable importance scores) for each target gene in order to identify the corresponding optimal list of TFs. We then re-iterate through the core GBM model using the optimal list of TFs for each target gene to obtain regularized transcriptional regulations followed by the additional refinement step in presence of any knockout information. The final inferred network obtained as a result of this procedure is directed and weighted.

Hereby, the proposed technique is referred as Regularized Gradient Boosting Machine (RGBM) for GRN inference. We evaluated RGBM on DREAM3, DREAM4 and DREAM5 network inference datasets and simulated RNA-Seq [24] datasets. The RGBM technique obtains superior performance in terms of higher values for *AU*_*pr*_ and *AU*_*roc*_ than the current state-of-the-art methods on majority of these datasets. Finally, we performed a case study by constructing the GRN for glioma tumors using gene expression profiles collected through the cancer genome atlas (TCGA) along with an apriori mechanistic ABN[1] of TFs and their corresponding targets with the purpose to identify the main regulators of the molecular subtypes of glioma.

## 2 MATERIALS AND METHODS

Several advantages of tree-based methods like GENIE [21], iRafNet [20] and ENNET [18] are that these approaches are computationally cheap, parallelizable, capture non-linear transcriptional regulations and are open-source. For in-silico networks of up to 100 genes in DREAM3 and DREAM4 challenges, these methods require less than a minute to infer the GRN. Here we propose a gradient boosting of regression stumps based framework (GBM) followed by a regularization step. This regularization step helps to identify the optimal set of TFs for each target gene and utilizes a series of transformations followed by a simple heuristic to detect and remove the targets which are not controlled by any regulator (i.e. upstream regulators or 0-indegree targets). Once the optimal set of TFs is obtained for each target, we re-iterate through the GBM framework to generate the edges. The resulting GRN is directed and the edges are directed from TF to target genes. The advantage of using GBM for GRN inference has been successfully showcased in [18].

A schematic representation of RGBM approach is illustrated in Figure 1. We first utilize a mechanistic active binding network (ABN) between TFs and their potential targets. The ABN is then fed as a *prior* information to the proposed machine learning framework. A Gradient Boosting Machine is used to to rank TFs that potentially regulate a target gene according to variable importance scores. The corner of L-curve shaped variable importance curve is used to identify the optimal set of TFs for a target gene. It also identifies upstream regulators by performing a series of transformation on the maximum variable importance score for each gene followed by using a simple heuristic cut-off. Finally we re-iterate through the boosting procedure with the optimal set of TFs for each target gene to assemble the final network. We describe the details of each step of Figure 1 in the following subsections.

**Figure 1:**
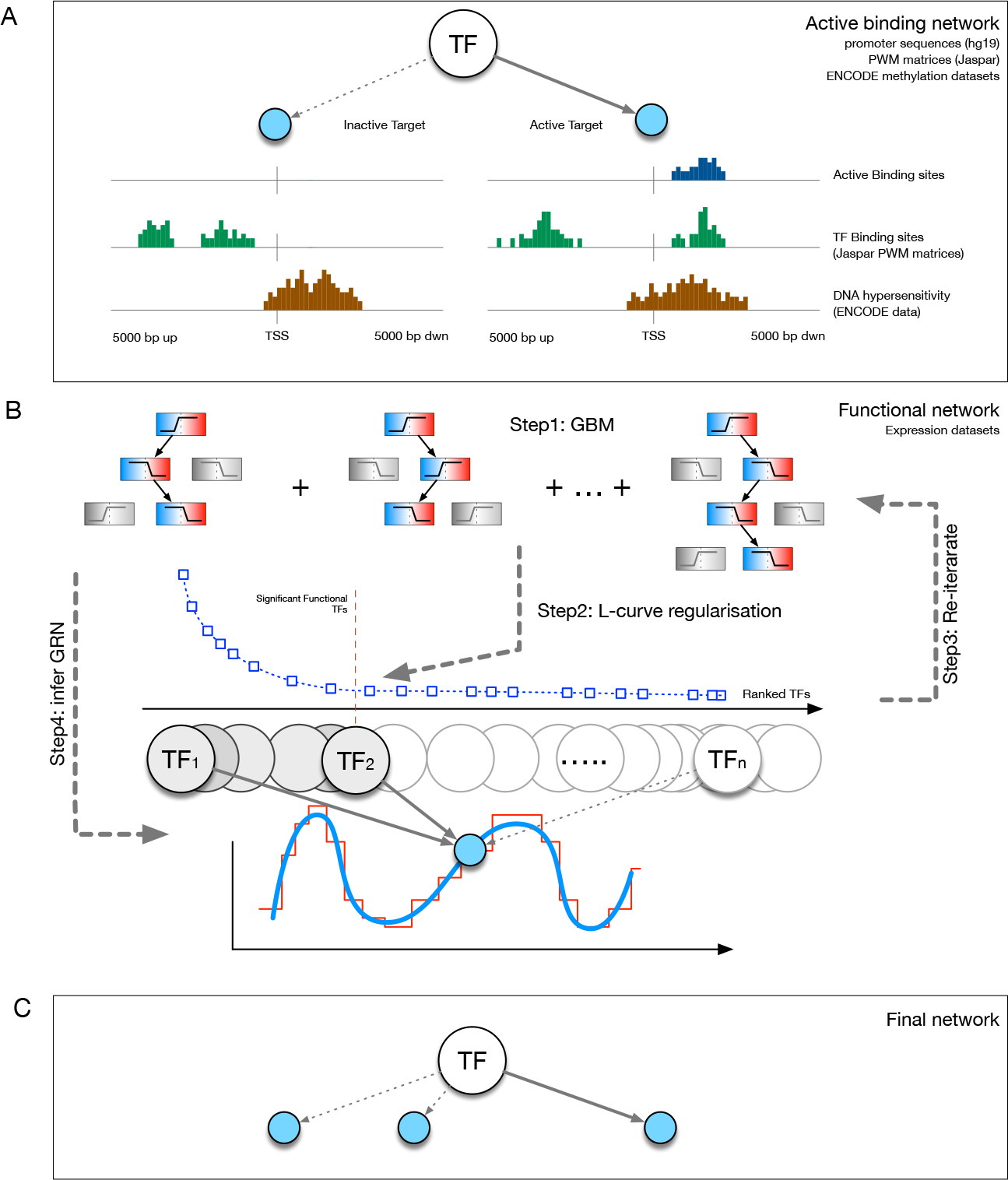
Schematic representation of the RGBM approach. (A) We first build the active binding network (ABN) and use it as *a priori* mechanistic network of connections between TFs and target genes. (B) illustration the primary procedure utilized by RGBM. Step1 uses a GBM to rank TFs that regulate a target gene according to variable importance scores. Step2 proposes a regularization step to locate the corner of L-curve shaped variable importance curve in order to identify the optimal set of TFs for a target gene. It also identifies upstream regulators by performing a series of transformation on the maximum variable importance score for each gene followed by using a simple heuristic cut-off. Step3 is to re-iterate through the boosting procedure with the optimal set of TFs for each target gene. Step4 is to infer the regulatory subgraph for each target gene. (C) representation the final inferred GRN obtained by combining the regulatory subgraphs of all target genes.

### 2.1 Pre-processing input data

An advantage of tree-based techniques is that they can take as input heterogeneous adjusted gene expression data together with any available meta-data information explaining the conditions of the experiments, for example, which genes were knocked out. In the case of the DREAM challenges, several additional information sources are available including knockout, knockdown and wildtype expression profiles. These describe the expression of a known set of perturbed genes, which are in a steady state at the time of measurement. On the other hand, time-series data describe the dynamics of expression levels of perturbed genes with no additional information, such as those available for knockout or knockdown experiments. The time-series data are usually extremely noisy and hence a smoothing step is essential to de-noise the data. We perform the smoothing of the time-series data using the “forecast” R package [25] for time-series data in the DREAM3 and DREAM4 challenges. We considered the fitted coefficients for each gene, for all the time steps/samples, as new smoothed noise-free version of time-series data. The variety of benckmark datasets used for GRN inference are illustrated in [18, 20, 26, 27]. Supplementary Figure 1 (F1) showcases two type of data used for GRNs.

An element of the expression matrix *E* ∈ 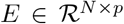 i.e. *e*_*ij*_, *i* = 1…*N* and *j* = 1…*p*, represents the expression value of the *j*^*th*^ gene in the *i*^*th*^ sample. The rows of *E* matrix correspond to the sample/experimental conditions and the columns represent the expression value of the genes in those samples (*p* genes in total). The additional (knockout or knockdown) information is maintained in a binary perturbation matrix 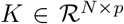 as in [18], where *K*_*ij*_ is a binary value corresponding to the *j*^*th*^ gene in the *i*^*th*^ experimental condition. If the *j*^*th*^ gene is perturbed in the *i*^*th*^ sample then *K*_*ij*_ = 1 otherwise *K*_*ij*_ = 0. In case of absence of any additional information, we set all values in the *K* matrix to 0.

### 2.2 Building the ABN

To learn potential regulatory activities between TFs and target genes, we merge constitutive associations due to active binding sites (ABN) and functional association due to contextual transcriptional activity (functional network) (Figure 1A).

The active binding network (ABN) is reconstructed from the collection of TF binding sites that are also active i.e. falling into not methylated regions. Binding sites are predicted with the FIMO (Find Individual Motif Occurrences) tool using 2, 532 unique motif PWMs (Position Weight Matrices) obtained from Jaspar [28] corresponding to 1, 203 unique TFs ([29, 30, 28, 31]). Instead, active promoter regions are classified with ChromHMM (v1.10), a Hidden Markov Model that classifies each genome position into 18 different chromatin states (9 states are considered open/active sites: *TssA, TssFlnkm, TssFlnkU, TssFlnkD, Tx, EnhG1, EnhG2, EnhA1m, EnhA1*) from 98 human epigenomes [32]. A binding relationship is considered active if the TF motif signal is significantly (FDR < 0.05) over-represented in the target promoter region (±5kbp TSS, hg19) and, in the same position (at least 1bp overlapping), the chromatin state is classified as open/active. The ABN consists of 6, 652, 518 overlapping active sites corresponding to 1, 959,125 unique TF associations between 1, 779 TFs and 51, 705 target genes.

### 2.3 From inference problem to a variable selection task

Given the rectified list of TFs i.e. *C*_*j*_ for each target gene *j* ∈ {1,…,*p*}, we sub-divide the problem of inferring the GRN into *p* independent tasks. For the *j*^*th*^ sub-problem, we get the subgraph corresponding to the outgoing edges from the appropriate TFs to the target gene *j*. To generate this subgraph, we first create an expression vector *Y*_*j*_ = *E*[, *j*] and a feature expression matrix *X*_*j*_ = *E*[,*C*_*j*_] from the expression matrix *E*. Here *Y*_*j*_ and *X*_*j*_ represent the dependent vector and the matrix of independent variables respectively. Supplementary Figure 2 (F2) illustrates how the *E* matrix is utilized to formulate *p* independent problems. Each of the *p* sub-problems can mathematically be formulated as:

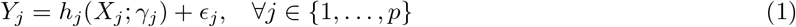

Here ϵ_*j*_ represents random noise and *h*_*j*_ (*X*_*j*_; *γ_j_*) is the parametric function that maps the expression matrix *X*_*j*_ to target (*Y*_*j*_) while optimizing the parameters *γ*_j_. Our goal is to identify a small number of TFs which drive the expression of the *j*^*th*^ gene using the columns of *X*_*j*_ as input features. Essentially, we have to solve a regression problem while inducing sparsity in the feature space resulting in a subset of the list of TFs (*SubC_j_*) which drive the expression of *j*^*th*^ target gene. This problem is also referred as feature or variable selection [33, 34] in statistical machine learning. Several statistical methods, including Lasso [35], Group Lasso [36], Fused Lasso [37], Elastic-net [38], perform a linear mapping from the feature space to the target space while inducing sparsity and have been utilized for GRN inference [8, 39, 40].

There are several tree-based machine learning methods, namely random-forests [41] and gradient boosting machines (GBM) [9], which solve the aforementioned problem using a non-linear mapping. Additionally, they provide a variable ranking scheme, which allows us to decipher directed edges from *SubC_j_* to the *j*^*th*^ target gene. These directed edges are weighted in accordance to the variable importance score that allows us to rank the variables. Tree-based methods have been extensively used for GRN reconstruction [21, 20, 18, 19, 42]. In this paper, we utilize a variant of the GBM model for inferring GRN. The working mechanism of the core GBM model is explained in detail in the Supplementary material. The authors in [18] utilized the LS-Boost (Supplementary Algorithm 1:S1) algorithm as the core GBM model to develop the ENNET [18] technique. In the proposed RGBM technique, we provide the user with the flexibility of utilizing either LS-Boost (Algorithm S1) or LAD-Boost (Algorithm S2) as core GBM model. This is because it was shown in [9] that LS-Boost performs extremely well for normally distributed expression values whereas LAD-Boost performs better for slash [9] distributed expression values.

Moreover, if knockout/knockdown expression data is available, we further apply a refinement step utilized in the null-mutant z-score method [7] and [18]. We reason that the direct regulation of a target by a TF would lead to a distinct signature in the expression data if the TF were knocked out. The details of the refinement step are also reported in the Supplementary material.

### 2.4 Main Regularization Steps

An important aspect of GRN is the sparsity, hence there are only a few TFs which regulate a target gene and there are a few genes which have no regulations i.e. we can have 0-indegree [4] in the inferred GRN. Thus, the procedure for reverse engineering GRN should return sparse networks and be able to detect such 0-indegree upstream regulators. From the initial set of steps (GBM + Refinement Step), the obtained adjacency matrix *A*_2_ can be quite dense as shown in Supplementary Figure 3 (F3).

However, when *A*_2_ is converted into an ordered edge-list (ranked in descending order based on edge-weights), several of the top ranked connections are indeed true, whereas many others can be false positives, as we showcase in our Results. But there is still a possibility to greatly reduce the number of falsely identified transcriptional regulations between the TFs and targets as showcased below.

We utilize a notion similar to the one used for optimizing the discrete L-curve criterion [22, 43] in Tikonov regularization [23]. This is done by analysing the variable importance (VI) curves (Figure 2) for individual target genes to obtain the optimal set of TFs for that gene. We also perform a series of transformations on the maximum VI (MVI) score for each target gene which is followed by a simple heuristic to detect 0-indegree genes in the final reverse-engineered GRN.

**Figure 2:**
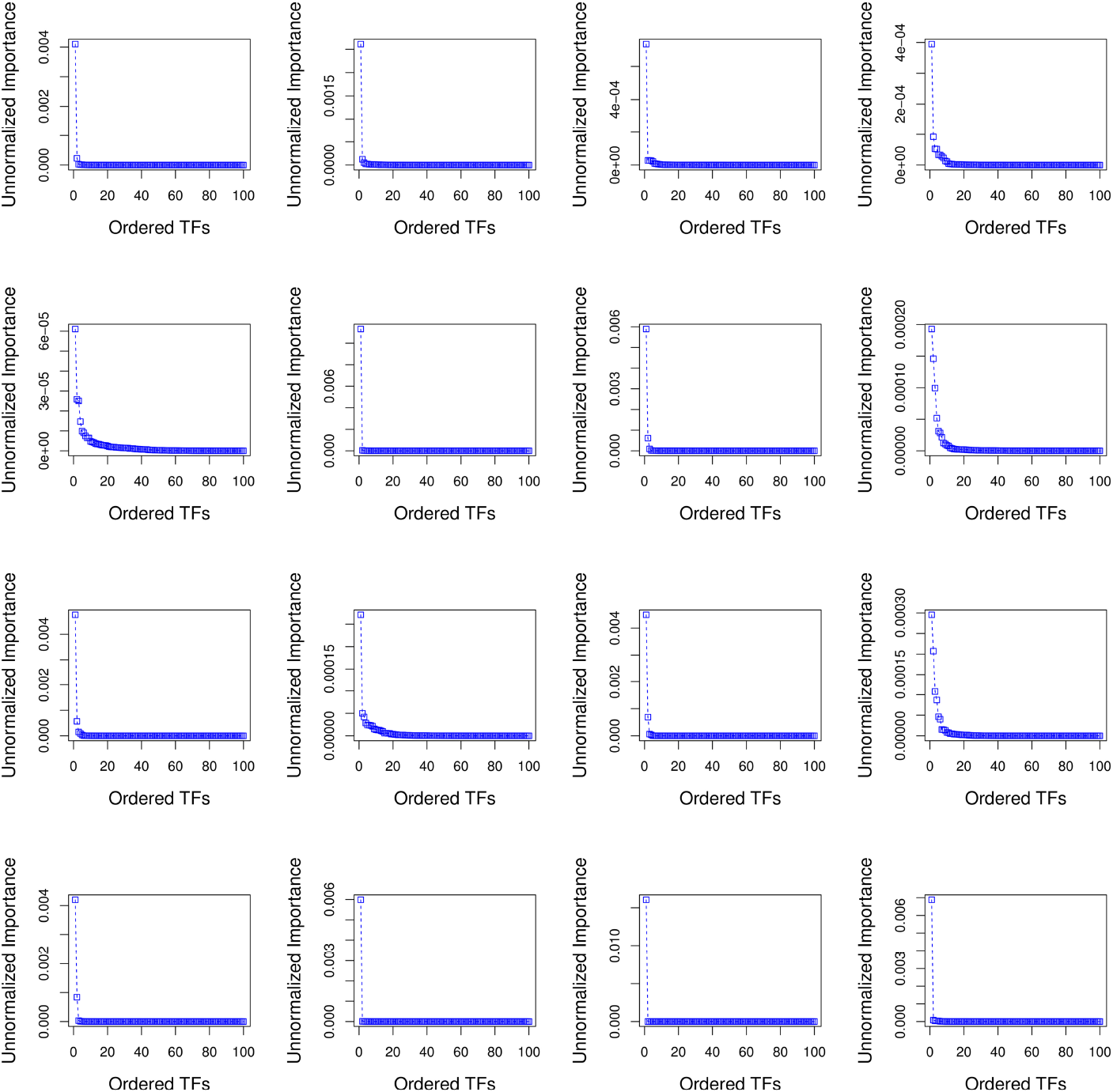
Sorted unnormalized variable importance (VI) curves for the first 16 targets of Network 1 from DREAM4 challenge. The target genes are ordered in row-format from left to right (i.e. 1 – 4 target genes in row 1,5 – 8 target genes in row 2, etc.) We can observe that the VI curves follow a Power-Law distribution w.r.t. the set of TFs for each target gene. This suggests that there are very few TFs which are strongly regulating the expression of individual target gene.

In order to identify the optimal set of TFs for each target gene, we use an idea similar to that used for identifying the corner in discrete L-curve criterion [43, 22] in Tikonov regularization [23]. The problem in Tikonov L-curve is to identify the corner of a discrete L-curve where the surface of the discrete L-curve is monotonically decreasing. Several algorithms have been proposed for computing the corner of a discrete L-curve, taking into account the need to capture the overall behaviour of the curve and avoiding the local corners [44, 45, 22]. In the case of sorted variable importance (VI) curve for individual target gene, the VI curve is also monotonically decreasing and usually follows a Power-Law distribution [46], as showcased in Figure 2. Moreover, rather than getting stuck in a local corner in the VI curve, we are inclined to locate the position where the VI curve first becomes flat w.r.t. the rightmost TF in the VI curve i.e. the position after which there is no change in the variable importance scores w.r.t. the smallest variable importance score in the VI curve. From Figure 2, we observe that the smallest variable importance score for a target gene is usually infinitesimal. The optimal set of TFs for the *j*^*th*^ target gene are those which are heavily regulating the expression of that target and hence *Imp_j_*(*ϕ*) > arg_min_(*Imp_j_*(*ϕ*)) for those TFs. To determine this optimal set of TFs for individual target gene, we use a modified variant of the *triangle method* [47] employed for identifying the corner in the Tikonov L-curve.

We first need to briefly explain the concept of the oriented angle between two line segments associated with a triple of points in the L-curve to explain the usage of the *triangle method* for identifying the optimal corner (optimal set of TFs) in the VI curve as shown in Figure 3. We consider the anti-clockwise convention to define an angle. Specifically, let 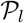, 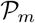 and 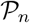 be three points on the VI curve satisfying *l < m < n* and let *v*_*m,l*_ denote the vector from 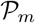 to 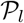. Then, we define the oriented angle *θ*(*l*,*m*,*n*) ∈ [−π, π] associated with the triplet as the angle between the two vectors *v*_*m,n*_ and *v*_*m,l*_ i.e. *θ*(*l*,*m*,*n*) = ∠(*v*_*m,n*_, *v*_*m*_,*l*). With this definition, an angle *θ*(*l*,*m*,*n*) = π corresponds to the point 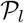, which determines the optimal position (optimal number of TFs) in VI curve. The key idea of the *triangle method* is to consider the following triples of L-curve points:

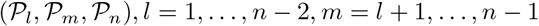

where *n* corresponds to the TF with the least non-zero importance score (*Imp_j_*(·)) for the *j*^*th*^ target gene. In the case of DREAM4 challenge (Network 1), *n* = *p* or the set of all genes. By using this idea, we identify as the corner the first triple where the oriented angle *θ*(*l*, *m*, *n*) is either equal to π or is maximum. If the angle *θ*(*l*, *m*, *n*) = π, then that part of the VI curve is already “flat” w.r.t. the least contributing TF and the position *l* represents the optimal number of TFs for the *j*^*th*^ target gene. All the TFs to the left of position *l* form the optimal set of TFs that regulate the target gene *j* as shown in Figure 3. The worst-case complexity of the *triangle method* is *O*(*p*^2^). However, since majority of the VI curves of the target genes have a Power-Law distribution, we can quickly reach the position where the oriented angle first becomes π and avoid unnecessary computations as indicated in Algorithm 1. From our experiments, we empirically found that the proposed technique requires *O(p)* steps to infer the optimal set of TFs for each target gene because of the Power-Law distribution of the VI scores. Moreover, the computation of the optimal set of TFs for each target gene can be performed independent of the other and hence is performed in parallel.

**Figure 3:**
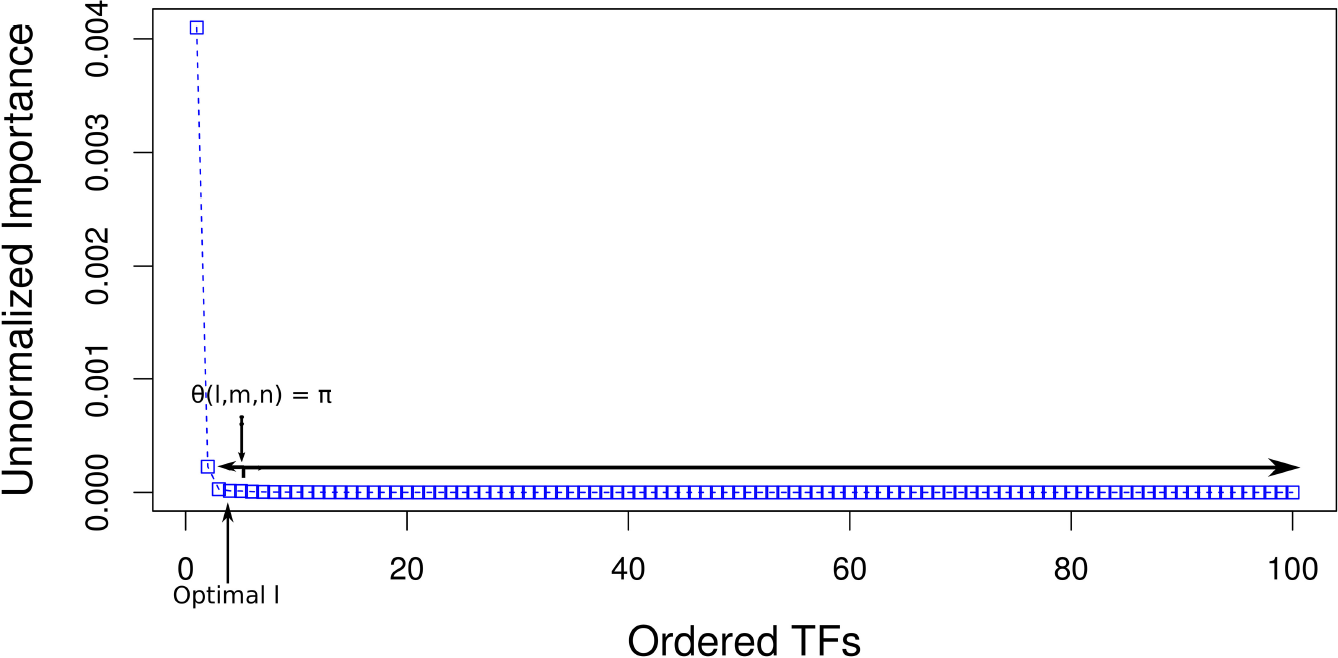
Optimal set of TFs obtained from the VI curve of gene “G1” for Network 1 from DREAM4 challenge using a *triangle method* [47] based technique which is commonly employed for identifying the corner in the Tikonov L-curve.

However, the computation of the optimal set of TFs for each gene is performed only after we have identified the upstream regulators i.e. 0-indegree nodes as explained earlier. From Figures 2 and 4A, we observe that the VI and MVI score distributions both follow Power-Law curves. From Figure 2, we observe that for several target genes the MVI score is *O*(10^−3^) or *O*(10^−4^). But there are a few outlier genes for which the MVI score is much smaller (*O*(10^−5^) or *O*(10^−6^)) indicating that the given set of TFs cannot drive the expression of these target genes. For example, for the 5^*th*^ target gene, the MVI score is ≈ 5 × 10^−5^ as shown in Figure 2.

Given the geometric nature of the distribution of MVI scores as observed from Figure 4A, it is difficult to directly select the scores which correspond to outlier targets i.e. genes which are not controlled by any TF. However, in case of a univariate Gaussian distribution, its relatively simple to define a heuristic cutoff (*ρ*) which helps to determine the outliers as indicated in Figure 4D. Hence, the problem of detecting 0-indegree genes can be solved efficiently by performing a set of transformations to convert a Power-Law distribution into a Gaussian distribution. It was shown in [48, 49] that given a univariate distribution, i.e. samples generated from a function of a single variable (*f*(*b*) = *e*^−λ*b*^ in our case), it can be turned into a normal distribution. In [49], the authors illustrated that the given distribution (*f*(*b*)) is first transformed to a uniform distribution and then an inverse of the cumulative density function of a normal distribution is applied on this uniform distribution to generate normally distributed values. Mathematically, it can be shown as:

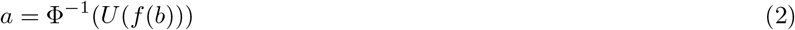

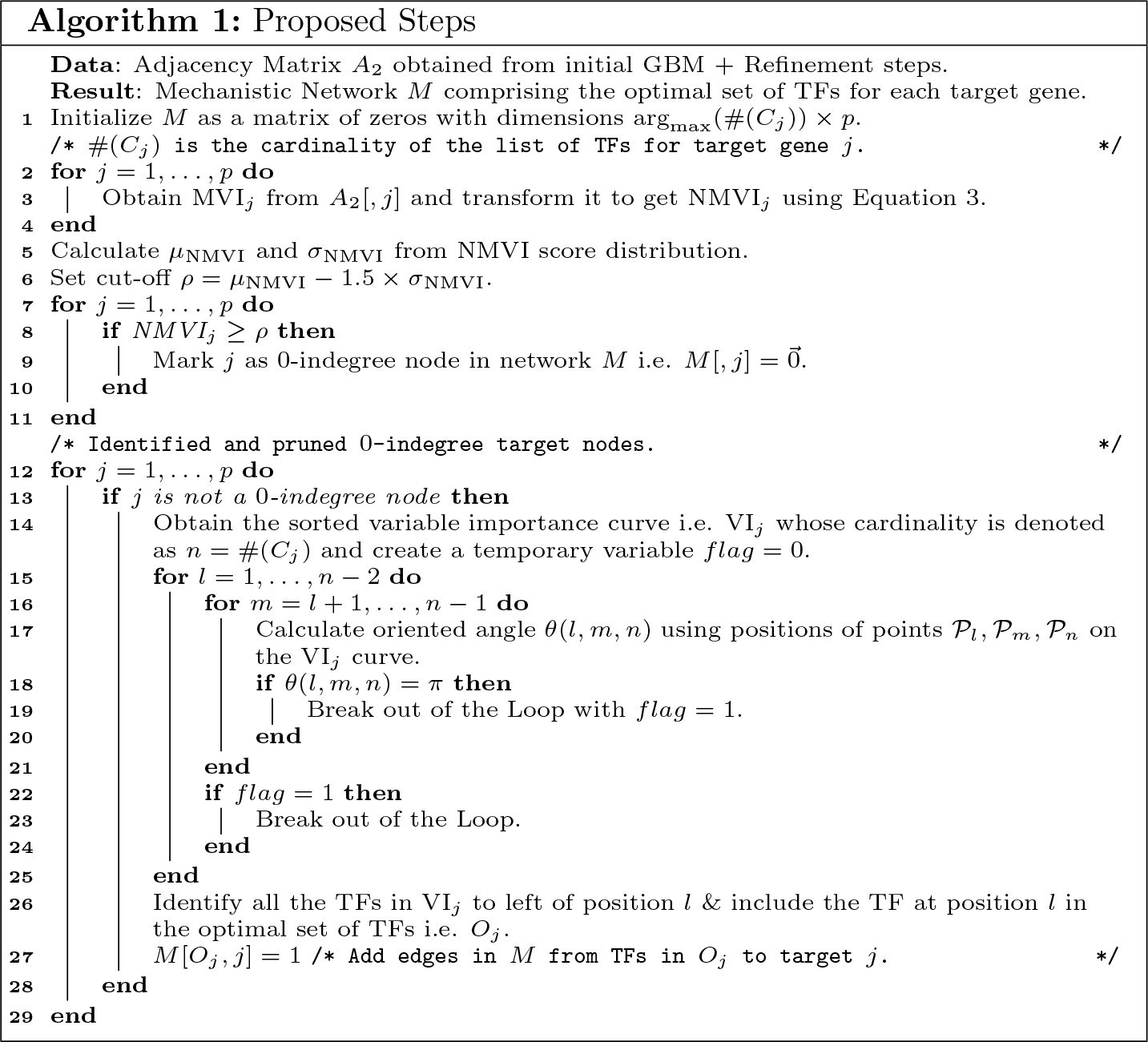

**Figure 4:**
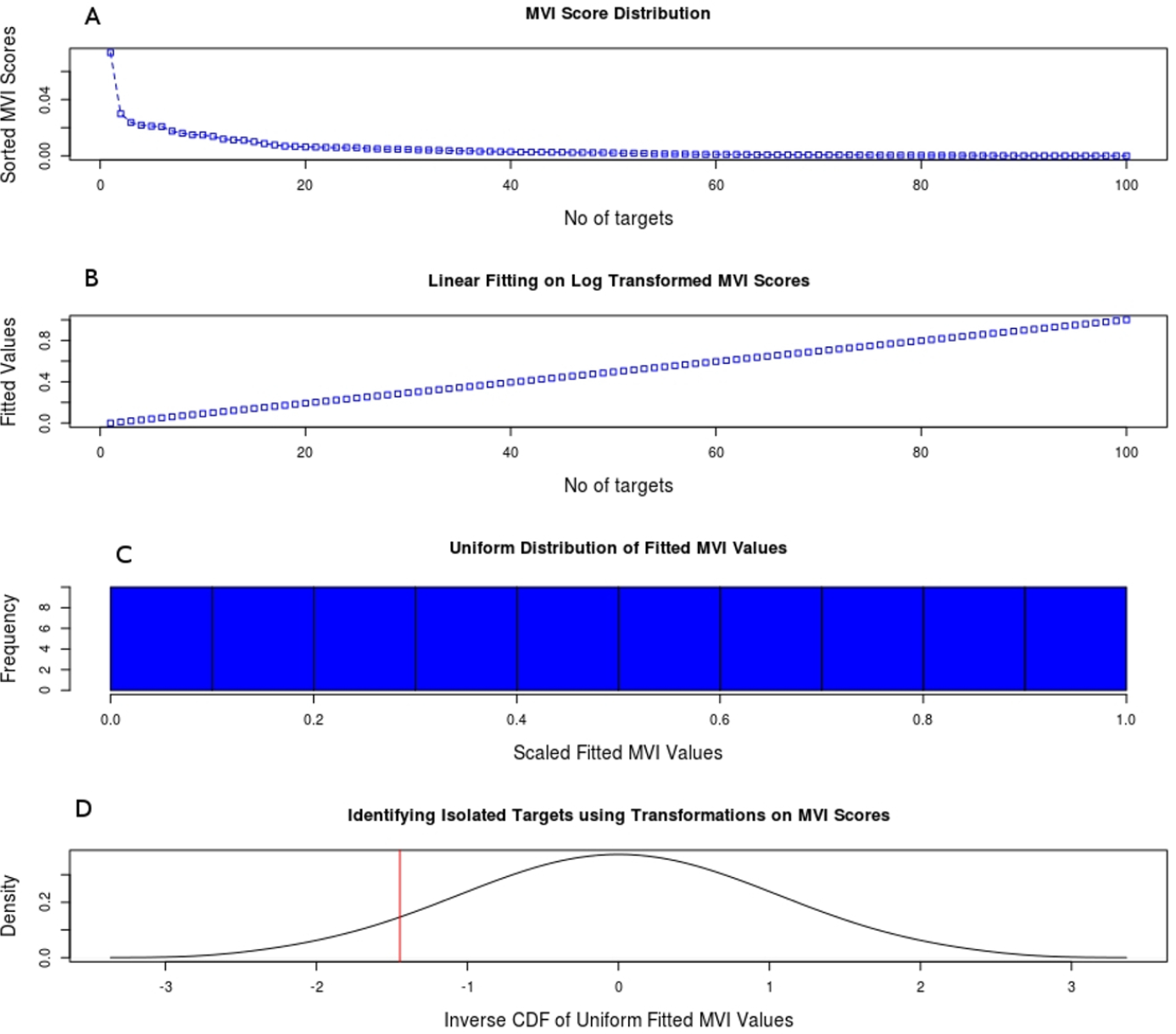
Subfigure A showcases that the MVI score distribution follows a Power-Law curve. Subfigure B represents the linear-fitting of log transformed MVI scores for all the 100 targets of Network 1 from DREAM4 challenge. Subfigure C illustrates that *U*(MVI_*j*_) = *S*(*lm*(log(MVI_*j*_))) follows a uniform distribution ∀*j* = 1,…,*p*. Subfigure D corresponds to the normally distributed MVI scores (NMVI). Here the “red” vertical line represents the heuristic cut-off selected to detect 0-indegree genes.

In the proposed framework, *f*(*b*) corresponds to the discrete distribution of MVI scores which follows a Power-Law curve as depicted in Figure 4A. The function *U*(·) transforms these MVI scores into uniformly distributed values in the range [0,1] as shown in Figure 4C. The function U(·) is a combination of a log(·) transformation on the initial MVI scores followed by a linear-fitting ((*lm*(·)) of the log(·) transformed values (Figure 4B). By performing this log(·) transformation and *lm*(·), smaller MVI scores end up getting larger negative values and we obtain a set of uniformly distributed values. However, these fitted values are still unscaled i.e. can take any values in ℝ. But it is necessary to scale (*S*(·)) these fitted values between the range [0,1] as it is essential for the inverse of the cumulative density function (Φ(·)) of a normal distribution i.e. Φ^−1^(·) in order to generate normally distributed scores (NMVI) as illustrated in Figure 4D. Here again smaller MVI scores correspond to smaller NMVI scores. These steps can mathematically be represented as:

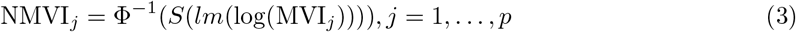

We obtain the mean (*μ*_NMVI_) and standard deviation (σ_NMVI_) values from the NMVI score distribution and select a heuristic cut-off *ρ* = μ_NMVI_−1.5*σ_NMVI_ to detect outlier NMVI scores (following the normally distributed nature of NMVI scores) i.e. NMVI_*j*_ ≤ *ρ* correspond to outlier target genes. These outlier scores correspond to the 0-indegree genes in the final inferred GRN (*A*_*final*_).

Once we have obtained the optimal set of TFs for an individual target gene (*M*), we re-iterate through the core GBM model using *M* as a mechanistic network and follow it with the refinement step. All these steps together form the RGBM technique for reconstructing GRNs as showcased in Algorithm 2 and illustrated via Figure 5.

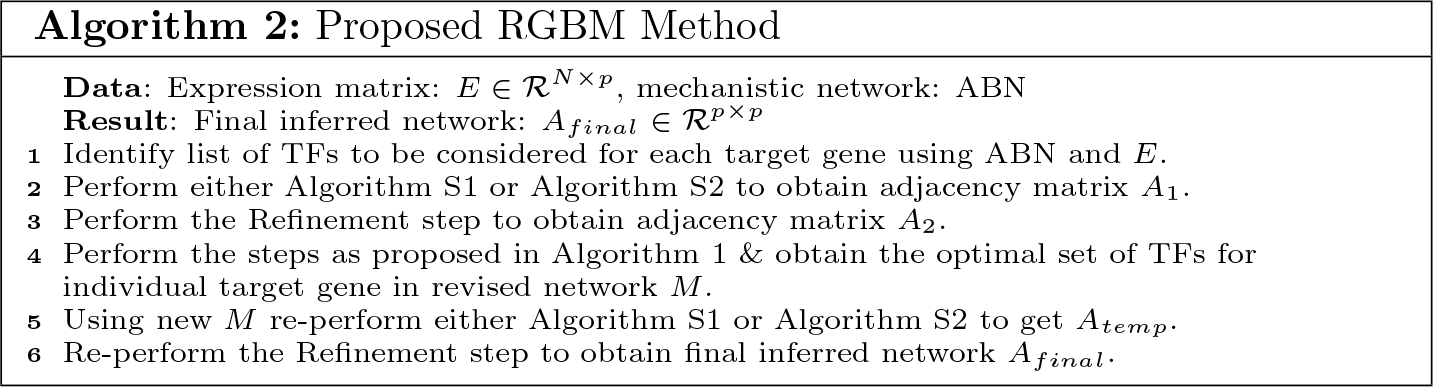

**Figure 5:**
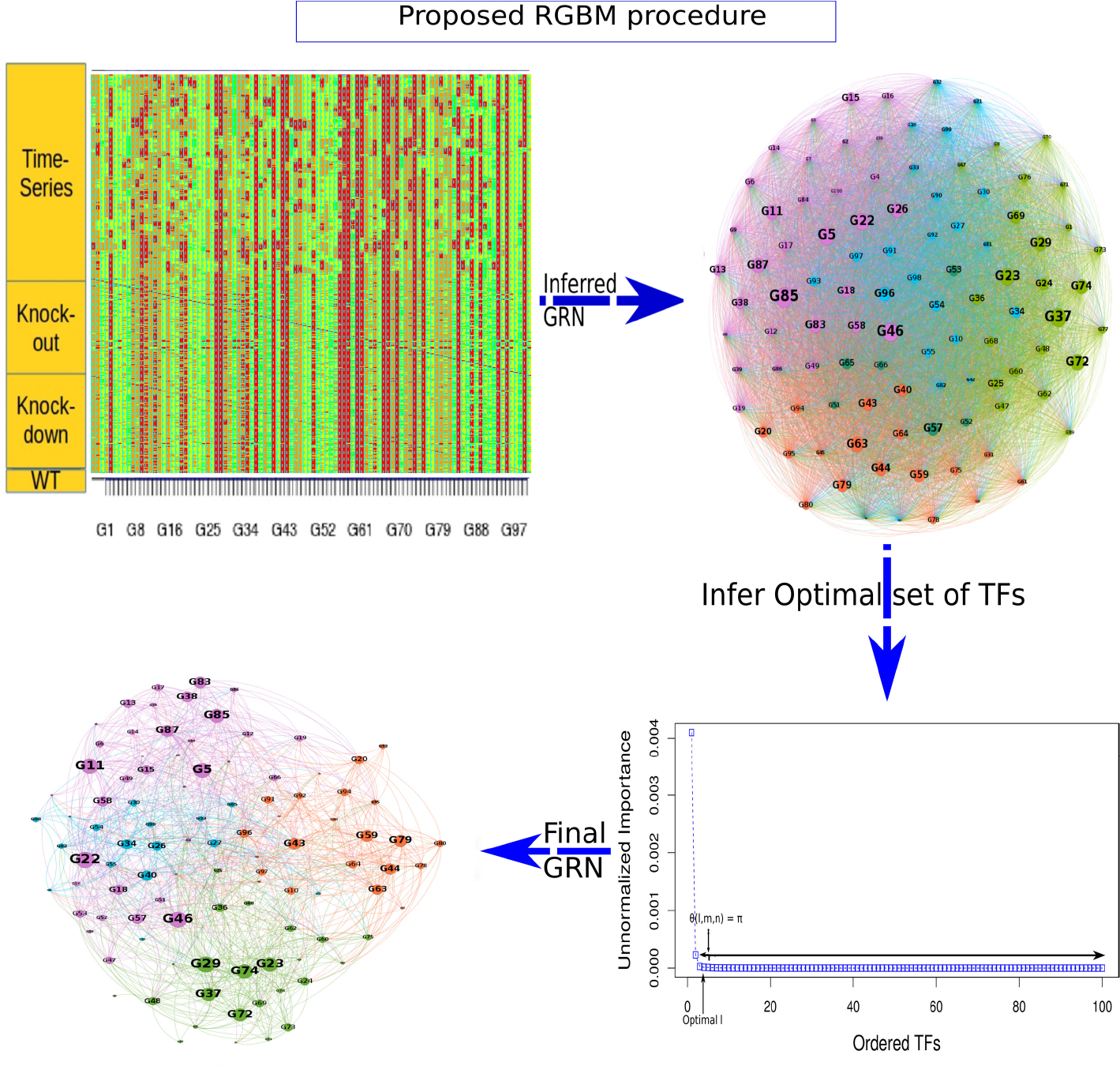
Final inferred GRN (*A*_*final*_) obtained as a result of RGBM Algorithm (Algorithm 2) on Network 1 for DREAM4 challenge. The final inferred GRN has 1,144 edges between 100 nodes whose edge weights are > 3.3 × 10^−15^ (machine precision). *A*_*final*_ is much more sparse in comparison to *A*_2_ which is obtained after initial GBM modelling followed by the Refinement step. In network *A*_*final*_, we have greatly reduced the number of falsely identified transcriptional regulations in *A*_2_.

### 2.5 Post-transcriptional TF activity

TF activity is determined using an algorithm that allows computationally inferring protein activity from gene expression profile data on an individual sample basis. The activity of a TF, defined as an index that quantifies the activation of the transcriptional program of a specific regulator in each sample *S*_*i*_, is calculated as follows:

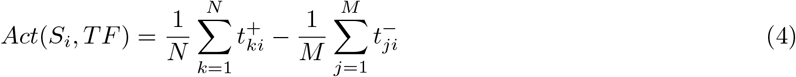

where 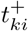 is the expression level of the *k*^*th*^ positive target of the MR in the *i*-th sample, 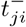 is the expression level of the *j*^*th*^ negative target of the MR in the *i*^*th*^ sample, *N* (*M*) the number of positive (negative) targets present in the regulon of the considered MR. If *Act*(*S*_*i*_,*TF*) > 0, the TF is *active* in that particular sample. f *Act* (*S*_*i*_, *TF*) < 0, the TF is inversely activated and if *Act* (*S*_*i*_, *TF*) ≈ 0 it is non-active. To identify the main Master Regulators of glioma subtypes reported in Section 3.4, we use supervised analysis on the activity function defined in (equation 4) using the Wilcoxon test [50].

## 3 RESULTS

Before we describe the experiments that we have undertaken to have a comprehensive comparison of the proposed method with other state-of-the-art GRN inference techniques, we briefly summarize the parameter settings for the RGBM models. The core of the RGBM model is the LS-Boost (Algorithm S1) or LAD-Boost (Algorithm S2) procedure (Algorithm 2, Step 1 and Step 4). The primary parameters of a GBM model are the number of iterations *T*, which is equivalent to the number of additive models built (Algorithms S1 and S2), the learning rate *v* and the sampling rate *s* for TFs during each iteration *t*. By default, we consider all the *N* samples/rows of the expression matrix *E* while building the GBM model. The authors of the ENNET procedure [18], whose underlying core model is also a GBM, and which currently is the state-of-the-art GBM model for reverse-engineering GRN, showcased through exhaustive experiments that the optimal set of values for the parameters of a GBM model is *T* = 5, 000, *v* = 0.001 and *s* = 0.3. We keep *T* = 5,000 as the upper bound on the number of iterations to perform boosting and utilize an early stopping criterion based on rate of change of average residue (*Ŷ*) values. We stop the boosting of regression stumps when this rate of change of average residue values between two iterations is below machine precision. Moreover, using these optimal set of values for the parameters of the GBM model, the network inferred as a result of the LS-Boost algorithm (i.e. Algorithm 2, Step 1) is equivalent to the one obtained from the ENNET technique. Hence, in all our experiments we used these optimal set of parameter values for building the core models (LS-Boost or LAD-Boost) in the proposed RGBM method.

### 3.1 RGBM outperforms state-of-the-art on DREAM Challenge Data

We assessed the performance of the proposed RGBM models using LS-Boost and LAD-Boost as core models on universally accepted benchmark networks of 100 or more genes from the DREAM3, DREAM4 and DREAM5 challenges [51, 52, 53] and compared them with several state-of-the-art GRN inference methods. For the purpose of comparison, we selected several methods including ENNET [18], GENIE3 [21], iRafNet [20], ARACNE [16] and the winner of each DREAM challenge. Among all the DREAM challenge networks, we performed experiments on the in-silico networks of size 100 from DREAM3 and DREAM4, and on 3 benchmark networks from DREAM5 challenge.

The DREAM3 challenge comprises 5 in-silico networks whose expression matrices *E* are simulated using GeneNetWeaver [54] software. Benchmark networks were constructed as sub-networks of systems of transcriptional regulations from known model organisms namely *E.coli* and *S.cerevisiae.* In our experiments, we focus on the networks of size 100, which are the largest in the DREAM3 suite. There are several additional sources of information available for these networks, such as knockout, knockdown and wildtype expressions apart from the time-series information. However, most of the state-of-the-art techniques do not necessarily utilize all these heterogeneous information sources. We showcase the best results generated for the DREAM3 challenge networks using the optimal combination of information sources for different GRN inference methods in Table 1. For all other combinations of information sources, the results of these GRN inference methods are inferior.

**Table 1:** Comparison of RGBM with myriad inference methods on DREAM3 networks of size 100. Here we provide the mean *AU*_*pr*_ and *AU*_*roc*_ values for 10 random runs of different inference methods. Here KO=Knockout, KD=Knockdown, WT=Wildtype and MTS=Modified smoothed version of the time-series data. The best results are highlighted in bold. *, +, − represent the quality metric values where RGBM (LAD-Boost), ENNET and iRafNet techniques respectively outperform the winner of DREAM3 challenge.

| Methods | Data Used | DREAM3 Experiments |  |  |  |  |  |  |  |  |  |
| --- | --- | --- | --- | --- | --- | --- | --- | --- | --- | --- | --- |
|  |  | Network 1 |  | Network 2 |  | Network 3 |  | Network 4 |  | Network 5 |  |
| | | $AU_{pr}$ | $AU_{roc}$ | $AU_{pr}$ | $AU_{roc}$ | $AU_{pr}$ | $AU_{roc}$ | $AU_{pr}$ | $AU_{roc}$ | $AU_{pr}$ | $AU_{roc}$ |
| RGBM (LS-Boost) | KO,KD,WT | **0.699** | 0.903 | **0.888** | **0.965** | **0.597** | 0.900 | **0.571** | **0.861** | **0.460** | **0.787** |
| RGBM (LAD-Boost) | KO,KD,WT | 0.683 | 0.903 | 0.870* | 0.963* | 0.562* | 0.900 | 0.535* | 0.853* | 0.400 | 0.770 |
| ENNET | KO,KD,WT,MTS | 0.627 | 0.901 | 0.865+ | 0.963+ | 0.552+ | 0.892 | 0.522+ | 0.842 | 0.384 | 0.765 |
| GENIE3 | KO,KD,WT | 0.430 | 0.850 | 0.782 | 0.883 | 0.372 | 0.729 | 0.423 | 0.724 | 0.314 | 0.656 |
| iRafNet | KO,KD,WT | 0.528 | 0.878 | 0.812- | 0.901 | 0.484 | 0.864 | 0.482- | 0.772 | 0.364 | 0.736 |
| ARACNE | KO,KD,WT | 0.348 | 0.781 | 0.656 | 0.813 | 0.285 | 0.669 | 0.396 | 0.662 | 0.274 | 0.583 |
| Winner [55] | KO, WT | 0.694 | **0.948** | 0.806 | 0.960 | 0.493 | **0.915** | 0.469 | 0.853 | 0.433 | 0.783 |

We observe from Table 1 that the best source of information for GENIE3, iRafNet and ARACNE methods are the knockout, knockdown and wildtype expressions. In presence of time-series data, the performance of these methods becomes worse. ARACNE performs the worst on DREAM3 challenge followed by tree-based GENIE3 and iRafNet techniques. ENNET method already defeats the winner [55] of DREAM3 challenge on several networks w.r.t. quality metric *AU*_*pr*_ and *AU*_*roc*_. However, RGBM using LS-Boost as core model significantly outperforms the winner on several networks of the DREAM3 challenge. This is because it gains maximum benefit from the proposed steps to remove falsely identified edges, efficiently detects 0-indegree genes and gains a lot in terms of precision and recall from the refinement step. The presence of additional knockout information greatly boosts the efficiency of the RGBM model. It was observed from the analysis of the later DREAM challenges [54] that the winning method for DREAM3 challenge made a strong assumption on the Gaussian type of measurement in the noise which was used in DREAM3 networks, but was no longer used in later DREAM challenges. As a result, the winning method of DREAM3 challenge was ranked 7^*th*^ in DREAM4 challenge.

The DREAM4 challenge took place one year after the DREAM3 challenge. It again comprised 5 benchmark networks which were constructed as sub-networks of system of transcriptional regulations from known model organisms namely *E.coli* and *S.cerevisiae*. We again focus on the networks of size 100 used in the DREAM4 suite. Additional sources of information including knockout, knockdown, wildtype and multifactorial information were also available for DREAM4 challenge. The best results generated for the DREAM4 challenge networks using the optimal combination of information sources for different GRN inference methods are depicted in Table 2. For all other combinations of information sources, the results of these GRN inference methods are inferior.

**Table 2:** Comparison of RGBM with myriad inference methods on DREAM4 networks of size 100. Here we provide the mean *AU*_*pr*_ and *AU*_*roc*_ values for 10 random runs of different inference methods. Here KO=Knockout, KD=Knockdown, WT=Wildtype and MTS=Modified smoothed version of the time-series data. The best results are highlighted in bold. *, + and − represent the quality metric values where RGBM, ENNET and iRafNet techniques outperform the winner of DREAM4 challenge.

| Methods | Data Used | DREAM4 Experiments |  |  |  |  |  |  |  |  |  |
| --- | --- | --- | --- | --- | --- | --- | --- | --- | --- | --- | --- |
|  |  | Network 1 |  | Network 2 |  | Network 3 |  | Network 4 |  | Network 5 |  |
| | | $AU_{pr}$ | $AU_{roc}$ | $AU_{pr}$ | $AU_{roc}$ | $AU_{pr}$ | $AU_{roc}$ | $AU_{pr}$ | $AU_{roc}$ | $AU_{pr}$ | $AU_{roc}$ |
| RGBM (LS-Boost) | KO,KD,WT,MTS | **0.709** | **0.936** | **0.561** | **0.878** | **0.525** | **0.911** | **0.616** | **0.903** | **0.450** | **0.893** |
| RGBM (LAD-Boost) | KO,KD,WT,MTS | 0.682<sup>*</sup> | 0.924<sup>*</sup> | 0.525<sup>*</sup> | **0.895** | 0.490<sup>*</sup> | 0.907<sup>*</sup> | 0.566<sup>*</sup> | **0.903** | 0.413<sup>*</sup> | 0.885<sup>*</sup> |
| ENNET | KO,KD,WT | 0.604<sup>+</sup> | 0.893 | 0.456<sup>+</sup> | 0.856<sup>+</sup> | 0.421<sup>+</sup> | 0.865<sup>+</sup> | 0.506<sup>+</sup> | 0.878<sup>+</sup> | 0.264<sup>+</sup> | 0.828<sup>+</sup> |
| GENIE3 | KO,WT | 0.338 | 0.864 | 0.309 | 0.748 | 0.277 | 0.782 | 0.267 | 0.808 | 0.114 | 0.720 |
| iRafNet | KO,TS | 0.552<sup>-</sup> | 0.901 | 0.337 | 0.799 | 0.414<sup>-</sup> | 0.835<sup>-</sup> | 0.421<sup>-</sup> | 0.847<sup>-</sup> | 0.298<sup>-</sup> | 0.792<sup>-</sup> |
| ARACNE | KO,KD,WT | 0.279 | 0.781 | 0.256 | 0.691 | 0.205 | 0.669 | 0.196 | 0.699 | 0.074 | 0.583 |
| Winner [56] | KO | 0.536 | 0.914 | 0.377 | 0.801 | 0.390 | 0.833 | 0.349 | 0.842 | 0.213 | 0.759 |

We observe from Table 2, that GBM based methods, RGBM (LS-Boost), RGBM (LAD-Boost) and ENNET clearly surpassed all the other methods including the winner of the DREAM4 challenge. The iRafNet and ENNET techniques also individually perform better than the winner [56] on several networks of DREAM4 challenge but they are easily outplayed by the two implementations of the RGBM method. RGBM method using the LS-Boost as core model has an exceptional performance on nearly all the networks w.r.t. the evaluation metrics in comparison to other state-of-the-art GRN reverse-engineering methods.

Figure 6 illustrates the optimal number of TFs identified by proposed Algorithm 1 for each target gene and passed as network *M* either to Algorithm S1 or Algorithm S2 followed by the refinement step to infer the final GRN for Network 1 of DREAM4 challenge. We observe that several genes (including “G5”, “G26”, “G40”, “G42” etc.) have 0 TFs connected to them and hence form the 0-indegree upstream regulators. Thus, in Figure 5 there are 0 incoming edges to these genes though there are some outgoing edges from these genes (for sub-problems where they act as TFs).

**Figure 6:**
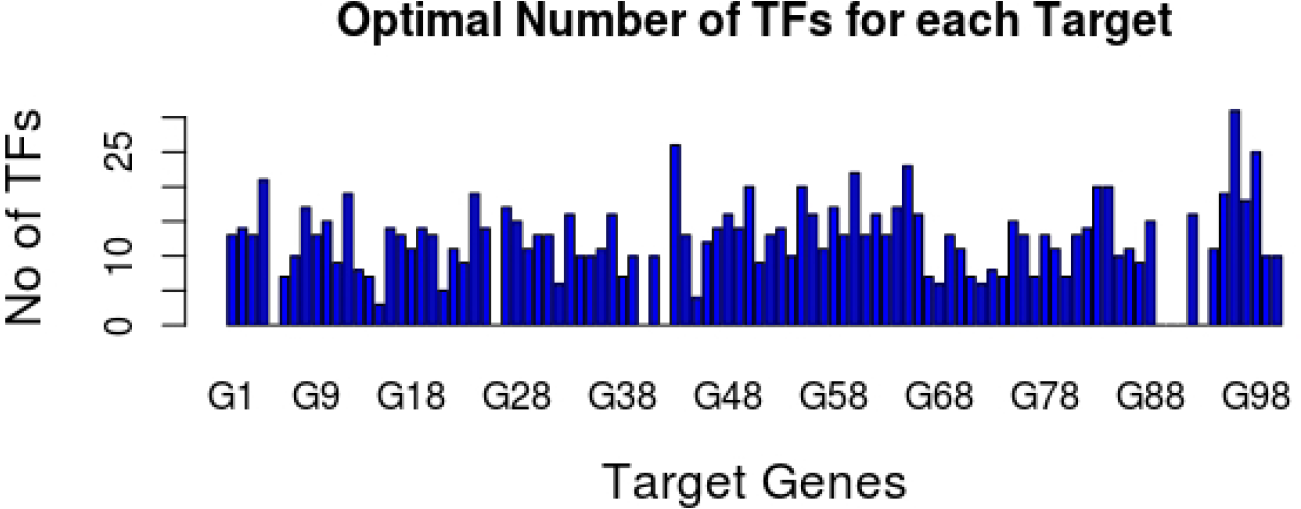
Optimal number of TFs for each target gene obtained from proposed Algorithm 1 for Network 1 of DREAM4 challenge.

Three benchmark networks in DREAM5 [4] challenge with different sizes and structure were generated using different model organisms. However, in this case, the time-series data of only one network was simulated in-silico, the two other sets of expression data were measured in real experiments in-vivo. As in previous DREAM challenges, in-silico expression data were simulated using the open-source GeneNetWeaver simulator [54]. However, DREAM5 was the first challenge where participants were asked to infer GRNs at for large-scale real datasets i.e. for thousands of target genes and hundreds of known TFs. Gold standard networks were obtained from two sources: RegulonDB database [57] and Gene Ontology (GO) annotations [58]. The results of all the inference methods for DREAM5 expression data using the optimal combination of information sources are summarized in Table 3.

**Table 3:** Comparison of RGBM with myriad inference methods on DREAM5 networks of varying sizes. Here we provide the mean *AU*_*pr*_ and *AU*_*roc*_ values for 10 random runs of different inference methods. Here KO=Knockout, KD=Knockdown, WT=Wildtype, MTS= Modified smoothed version of the time-series data, Exp=Steady-state gene expression. The best results are highlighted in bold. *, + and − represent the quality metric values where RGBM, ENNET and iRafNet techniques respectively defeat the winner of DREAM5 challenge i.e. GENIE3 method.

| Methods | Data Used | DREAM5 Experiments |  |  |  |  |  |
| --- | --- | --- | --- | --- | --- | --- | --- |
|  |  | Network 1 |  | Network 3 |  | Network 4 |  |
| | | $AU_{pr}$ | $AU_{roc}$ | $AU_{pr}$ | $AU_{roc}$ | $AU_{pr}$ | $AU_{roc}$ |
| RGBM (LS-Boost) | KO,Exp | **0.537** | 0.846* | 0.086 | 0.633* | **0.048** | **0.546** |
| RGBM (LAD-Boost) | KO,Exp | 0.513* | 0.842* | 0.084 | 0.628* | 0.047* | 0.544* |
| ENNET | KO,Exp | 0.432<sup>+</sup> | **0.857** | 0.069 | 0.632<sup>+</sup> | 0.021 | 0.532<sup>+</sup> |
| iRafNet | KO,MTS,Exp | 0.364<sup>-</sup> | 0.813 | **0.112** | **0.641** | 0.021 | 0.523<sup>-</sup> |
| GENIE3 (Winner) | Exp | 0.291 | 0.814 | 0.094 | 0.619 | 0.021 | 0.517 |
| TIGRESS [8] | KO,Exp | 0.301 | 0.782 | 0.069 | 0.595 | 0.020 | 0.517 |
| CLR [15] | Exp | 0.217 | 0.666 | 0.050 | 0.538 | 0.018 | 0.505 |
| ARACNE | Exp | 0.099 | 0.545 | 0.029 | 0.512 | 0.017 | 0.500 |

RGBM using LS-Boost core model gives better results than other methods w.r.t evaluation metrics *AU*_*pr*_ and *AU*_*roc*_ on Network 4 as illustrated in Table 3. It easily defeats the winner (GENIE3) of DREAM5 challenge and outperforms recent state-of-the-art GRN inference methods iRafNet and ENNET techniques. However, the predictions for in-vivo expression profiles (DREAM5 challenge - Network 3 and Network 4) result in extremely low precision-recall values as depicted in Table 3. One of the reasons for the poor performance of all the inference methods for such expression data is the fact that experimentally derived pathways, and consequently gold standards obtained from them, are not necessarily complete, regardless of how well the model organism is known. Additionally, there are regulators of gene expression other than TFs, such as miRNA and siRNA, which also drive the expression of these genes. As shown in this study, an in-silico expression matrix provide enough information to confidently reverse-engineer their underlying structure, whereas in-vivo data hide a much more complex system of regulatory interactions.

### 3.2 RGBM outperforms state-of-the-art on Synthetic RNA-Seq Data

We conducted additional experiments on simulated RNA-Seq data. We used our “synRNASeqNet” [59] package in **R** to generate 5 RNA-Seq expression matrices using the “simulatedData” function in the package. The “simulatedData” function uses a stochastic Barabási-Albert (BA) model [46] to build random scale-free networks using a preferential attachment mechanism with power exponent *α* and simulated RNA-Seq counts from a Poisson multivariate distribution [60]. For our experiments, we generated 5 RNA-Seq expression (*E*) matrices comprised of 500 RNA-Seq counts for 50 target genes using power exponent values *α* ∈ {1.75, 2, 2.25, 2.5, 2.75} respectively. For each experiment, we are not provided with any additional information (absence of perturbation matrix *K*), and for each target gene we consider all the remaining genes as TFs while reverse-engineering the GRN. We again used evaluation metrics like *AU*_*pr*_ and *AU*_*roc*_ to compare the proposed RGBM (using LS-Boost as core model) with state-of-the-art GRN inference methods, including ENNET, GENIE3 and ARACNE. Supplementary Figure 4 (F4) illustrates the comparison of various GRN inference methods w.r.t. the Receiver Operating Characteristics (ROC) for each of the five experiments. Similarly, Supplementary Figure 5 (F5) showcases the precision-recall curves, used to estimate the *AU*_*pr*_ metric, for different GRN inference methods.

The performance of RGBM is compared with ENNET, GENIE3 and ARACNE for five different experimental settings as shown in Table 4. For each of these experiments, we increased the value of the preferential attachment parameter *α* in the BA-model from 1.75 to 2.75 in steps of 0.25, thereby making the task of re-constructing accurate GRNs from simulated RNA-Seq counts harder as observed from the *AU*_*roc*_ and *AU*_*pr*_ values in Table 4.

**Table 4:** Comparison of proposed RGBM technique with ENNET, GENIE and ARACNE GRN inference methods w.r.t. evaluation metrics *AU*_*roc*_ and *AU*_*pr*_ for reverse-engineering GRNs from RNA-Seq counts where the underlying ground-truth network follows a BA preferential attachment model with exponent *α*.

| Methods | RNA-Seq Experiments |  |  |  |  |  |  |  |  |  |
| --- | --- | --- | --- | --- | --- | --- | --- | --- | --- | --- |
| | Exponent $\alpha = 1.75$ | | Exponent $\alpha = 2$ | | Exponent $\alpha = 2.25$ | | Exponent 2.5 | | Exponent 2.75 | |
| | $AU_{pr}$ | $AU_{roc}$ | $AU_{pr}$ | $AU_{roc}$ | $AU_{pr}$ | $AU_{roc}$ | $AU_{pr}$ | $AU_{roc}$ | $AU_{pr}$ | $AU_{roc}$ |
| RGBM | 0.561 | 0.802 | 0.448 | 0.781 | **0.522** | **0.686** | **0.521** | **0.686** | **0.520** | **0.686** |
| ENNET | 0.554 | 0.800 | **0.452** | **0.785** | 0.498 | 0.680 | 0.498 | 0.680 | 0.498 | 0.679 |
| GENIE3 | **0.566** | **0.854** | 0.439 | 0.782 | 0.195 | 0.681 | 0.184 | 0.676 | 0.176 | 0.669 |
| ARACNE | 0.065 | 0.588 | 0.054 | 0.568 | 0.056 | 0.623 | 0.055 | 0.612 | 0.055 | 0.612 |

Here the evaluation metrics *AU*_*roc*_ and *AU*_*pr*_ represents the mean value of these evaluation metrics for 10 random runs of each setting. We can observe from Table 4 that the RGBM and ENNET methods are robust w.r.t. increasing power exponent *α* used for generating the BA model [46] based networks. This is because the performance of these methods is not affected by increasing values of α whereas the effectiveness of GENIE3 and ARACNE GRN inference methods decreases with increase in α. In particular, the performance of GENIE3 decreases drastically w.r.t. the evaluation metric AUpr for increasing values of preferential attachment exponent α. The RGBM technique outperforms other GRN inference methods on 3 out of the five synthetic networks as illustrated in Table 4 and Figures F4 and F5, thereby showcasing its superiority.

### 3.3 Evaluation of RGBM on E.Coli SOS, S.Cerevisiae Cell Cycle and IRMA data

We compared the performance of RGBM on classical real datasets with a known underlying network. We used the following datasets:

- *E.coli* SOS pathway. Eight genes regulating the SOS response of DNA damage. Following our previous work [17], we choose genes involved in DNA damage tolerance and repair activated through recA and lexA. We adopt the first 14 point time course profiles of an experiment made publicly available by Ronen et al. [61].
- *S.cerevisiae* cell cycle. An eleven gene network controlling the G1 step of cell cycle. As in [17], we choose genes whose mRNA levels respond to the induction of Cln3 and Clb28, two cell cycle regulators, and a 16 point time-course profiles made publicly available by Spellman et al. [62]
- *IRMA*. An in-vivo fully controlled experimental network built in *S.cerevisiae* and composed of five genes [63]. The network is perturbed by culturing cells in the presence of galactose or glucose. Galactose activates the GAL1-10 promoter, cloned upstream of a Swi5 mutant in the network, and it is able to activate the transcription of all the five network genes. Two perturbations corresponding to growing cells from glucose to galactose medium and to reverse shift are made respectively for the 16 and 21 time points.

Since the above datasets correspond to time-course experiments, we include in Table 5 the results on three methods specifically developed for time-course data: TimeDelay-Aracne [17], FormalM [64] and TSNIF [65]. We can see from Table 5 that RGBM outperforms other approaches in terms of F1 metric for the *E. Coli* and IRMA networks, whereas it reaches the best recall for the *S.cerevisiae* cell cycle.

**Table 5:**
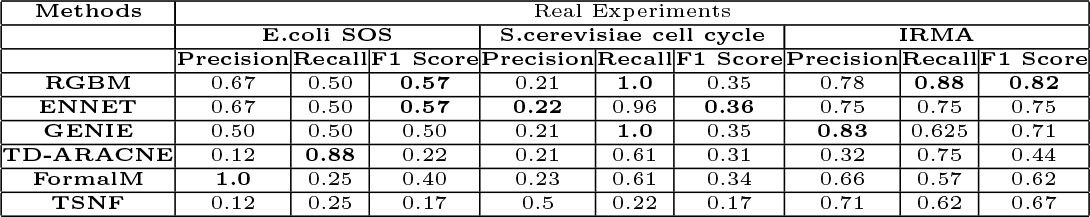
Comparison of RGBM with ENNET, GENIE, Time-Delay ARACNE, FormalM, TSNF, Banjo GRN inference methods w.r.t. evaluation metrics Precision, Recall and F1 Score for reverse-engineering GRNs from E.coli SOS, S.cerevisiae cell cycle and IRMA expression data. The best results are highlighted in bold.

### 3.4 RGBM Identifies the Master Regulators of Glioma Cancer Subtypes

The results in the previous paragraphs have shown that RGBM is able to efficiently recover the regulatory structure of small and large gene networks. Here we apply RGBM for the identification of Master Regulators of tumor subtypes of human glioma, the most frequent primary brain tumor in adults [66]. In the cancer field, Master Regulators (MR) have been defined as gene products (mostly TFs) necessary and sufficient for the expression of particular tumor-specific signatures typically associated with specific tumor phenotypes (e.g. proneural vs. mesenchymal). In the case of malignant gliomas, reverse engineering was used to successfully predict the experimentally validated transcriptional regulatory network responsible for activation of the highly aggressive mesenchymal gene expression signature of malignant glioma [67]. Therefore, a Master Regulator gene can be defined as a network hub whose regulon exhibits a statistically significant enrichment of the given phenotype signature which expresses a cellular phenotype of interest such as tumor subtype. MARINa (MAster Regulator INference algorithm) is an algorithm to identify MRs starting from a GRN and a list of differentially expressed genes [68]. This specific algorithm was succesully applied to identify Stat3 and C/EBPβ as the two TFs hierarchically placed at the top of the transcriptional network of mesenchymal high-grade glioma [67].

Recently, we worked with the Analysis Working Group (AWG) of the The Cancer Genome Atlas-TCGA project to deconvolute a PanGlioma dataset that included the largest collection of human glioma ever reported [69]. We reported that, using a combination of DNA copy number and mutation information, DNA methylation and mRNA gene expression, human gliomas can be robustly divided into seven major subtypes that we defined as G-GIMP-low, G-CIMP-high, Codel, Mesenchymal-Like, Classic-Like, LGm6-GBM and PA-like [69]. The first key division of human glioma is driven by the status of the IDH1 gene, whereby IDH1 mutations are typically characterized by a relatively more favorable clinical course of the disease. IDH1 mutation are associated with an hypermethylation phenotype of glioma (G-CIMP, [70]). However, our PanGlioma study reported that IDH-mutant tumors lacking codeletion of chromosome 1p and 19q is an heterogeneous subgroup characterized by a predominant G-CIMP-high subtype and a less frequent the G-CIMP-low subgroup that is characterized by relative loss of the DNA hypermethylation profile, worse clinical outcome and likely reporesents the progressive evolution of G-CIMP-high gliomas towards a more aggressive tumor phenotype [69]. However, the transcriptional network and the set of MRs responsible for the transformation of G-CIMP-high into G-CIMP-low gliomas remained elusive.

Among the large group of IDH-wildtype tumors (typically characterized by a worse prognosis when compared to IDH-mutant glioma), we discovered that, within a particular methylation-driven cluster (LGm6) and at variance with the other methylation-driven clusters of IDH-wildtype tumors, the lower grade gliomas (LGG) display significantly better clinical outcome than GBM tumors (GBM-LGm6). We defined these LGG tumros as PA-like based on their expression and genomic similarity with the pediatric tumor Pylocitic Astrocytoma. However, as for the transition from G-CIMP-high into G-CIMP-low gliomas, the determinants of the malignant progression of PA-like LGG into GBM-LGm6 remained unknown. Here, we applied our novel computational RGBM approach to infer the MRs responsible for the progression of G-CIMP-high into G-CIMP-low IDH-mutant glioma and those driving progression of PA-like LGG into LGm6-GBM IDH-wildtype tumors respectively.

Towards this aim, we first built the PanGlioma network between 457 TFs and 12,985 target genes. An ABN network was to used as prior for the RGBM algorithm and for the expression matrix we used the TCGA pan-glioma dataset [69] including 1250 samples (463 IDH-mutant and 653 IDH-wild-type), 583 of which were profiled with Agilent and 667 with RNA-Seq Illumina HiSeq downloaded from the TCGA portal. The batch effects between the two platforms were corrected as reported in [71] using the COMBAT algorithm [72] having tumor type and profiling platform as covariates. Subsequently, quantile normalization is applied to the whole matrix. The network is shown in supplementary Figure 6 (F6) and contains 39,192 connections with an average regulon size of 85.8 genes. To identify the MRs displaying the highest differential activity for each group, we ranked MR activity for each TF among for all the 7 glioma subtypes. We also represent in the Extended Figures 8-14 how each MR is activated in the various subtypes.

The top MRs exhibiting differential activity among the glioma groups are shown in supplementary Figure 7 (F7) and their average activity in Figure 7. We found that RGBM-based MR analysis efficiently separates an IDH-mutant dominated cluster of gliomas including each of the 3 IDH-mutant subtypes (G-CIMP-high, G-GIMP-low, and Codel) from an IDH-wildtype group including Mesenchymal-Like, Classic-Like and LGm6-GBM. This finding indicates that RGBM correctly identifies biologically-defined subgroups in terms of the activity of MRs. The MRs characterizing IDH-mutant glioma include known regulators of cell fate and differentiation of the nervous system, therefore, indicating that these tumors are driven by a more differentiated set of TFs that are retained from the neural tissue of origin (e.g. NEUROD2, MEF2C, EMX1, etc.). Conversely, the MRs whose activity is enriched in IDH-wildtype glioma are well-known TFs driving the mesenchymal transformation, immune response and the higher aggressiveness that characterizes IDH-wildtype glioma (STAT3, CEBPB, FOSL2, BATF and RUNX2, etc). Remarkably, while the G-CIMP-low subtype showed a general pattern of activation of MRs that includes this subtype within the IDH-mutant group of gliomas, when compared to the G-CIMP-high subtype, G-CIMP-low glioma displays distinct loss of activation of neural cell fate/differentiation-specific MRs (see for example the activity of the crucial neural TFs NEUROD2, MEF2C and EMX1) with corresponding activation of a small but distinct set of TFs that drive cell cycle progression and proliferation (E2F1, E2F2, E2F7 and FOXM1). This finding indicates that the evolution of the G-CIMP-high into the G-CIMP-low subtype of glioma is driven by (i) loss of the activity of neural-specific TFs and (ii) gain of a proliferative capacity driven by activation of cell cycle/proliferation-specific MRs.

**Figure 7:**
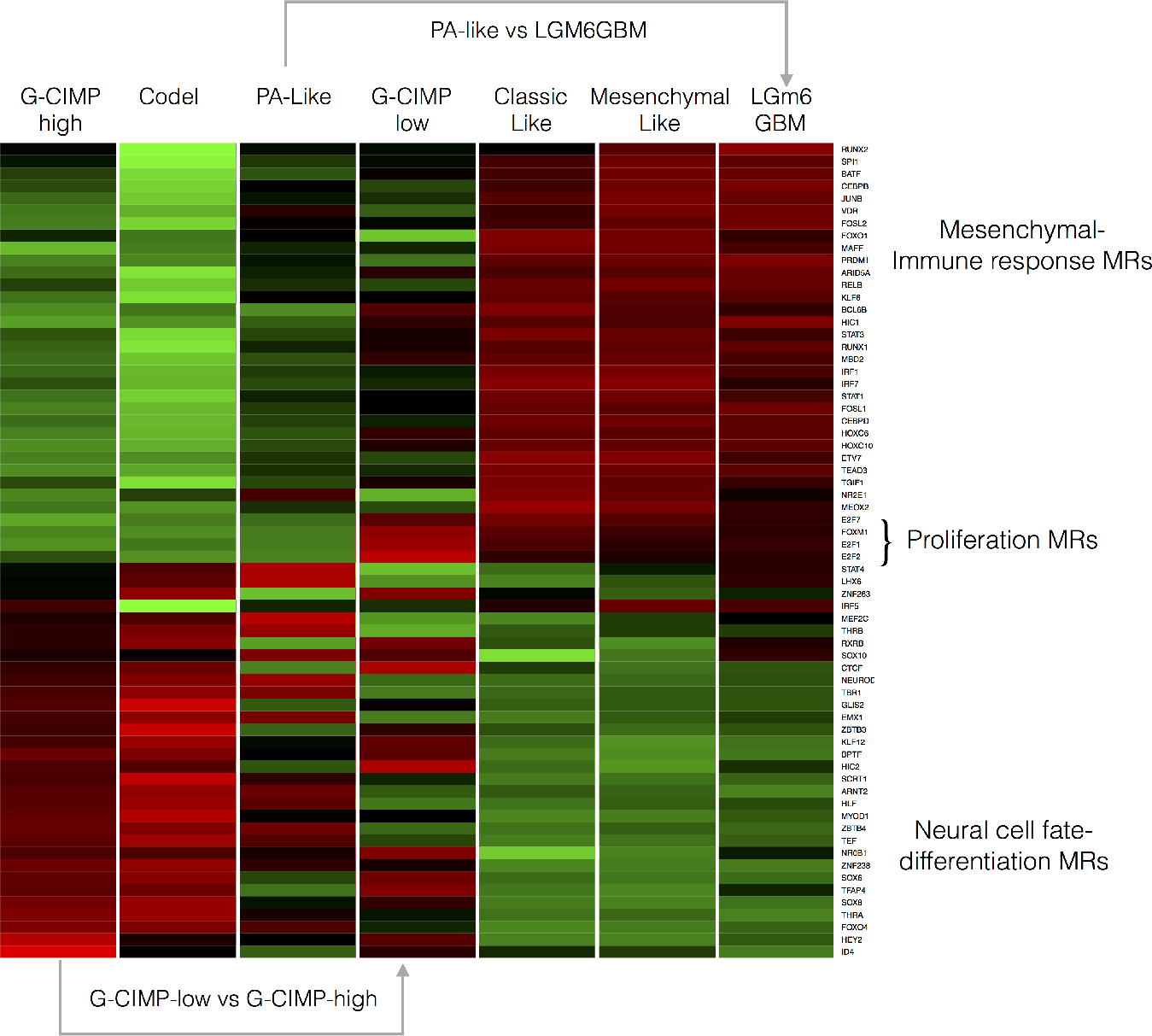
Average MR activity in the seven glioma subtypes.

Concerning the PA-like into LGm6-GBM, we note that, despite being sustained by an IDH-wildtype status, PA-like LGG cluster within the IDH-mutant subgroup of glioma, with higher activity of Neural cell fate/differentiation-specific MRs and inactive Mesenchymal-immune response MRs. Therefore, the evolution of PA-like LGG into LGm6-GBM is marked by gain of the hallmark aggressive MR activity of high grade glioma with corresponding loss of the MRs defining the neural cell of origin of these tumors.

Taken together, the application of the RGBM approach to the recently reported Pan-Glioma dataset revealed the identity and corresponding biological activities of the MRs driving transformation of the G-CIMP-high into the G-CIMP-low subtype of glioma and PA-like into LGm6-GBM, thus, providing a clue to the yet undetermined nature of the transcriptional events driving the evolution among these novel glioma subtypes.

## 4 CONCLUSION

In this paper, we proposed a novel GRN inference method namely, RGBM, whose core model for deducing transcriptional regulations for each target gene is boosting of regression stumps. We showcased that RGBM provides efficient results with both the LS-Boost and the LAD-Boost loss functions. Several contributions of RGBM are a) incorporation of prior knowledge in the form of a mechanistic ABN i.e. to consider previously known regulations between TF-targets; b) consideration of multiple heterogeneous sources of information to build the expression profiles i.e. time-series, knockout, knockdown and wild-type information; c) utilization of the idea that a given target gene is regulated by only a few TFs and proposed a novel technique based on **triangle method** [47], employed for identifying the corner in the Tikonov L-curve, to identify the optimal set of TFs for each target gene; d) identification of several upstream regulators as 0-indegree nodes in the final inferred GRN. We performed a thorough evaluation of RGBM on DREAM challenge datasets and simulated RNA-Seq datasets w.r.t. evaluation metrics *AU*_*roc*_ and *AU*_*pr*_ and demonstrated that RGBM easily defeats other GRN inference techniques including ENNET, GENIE3 and ARACNE in these experiments.

## References

[1] Christopher L Plaisier, Sofie OBrien, Brady Bernard, Sheila Reynolds, Zac Simon, Chad M Toledo, Yu Ding, David J Reiss, Patrick J Paddison, and Nitin S Baliga. Causal mechanistic regulatory network for glioblastoma deciphered using systems genetics network analysis. Cell Systems, 3(2):172–186, 2016.

2 EP van Someren, LFA Wessels, Eric Backer, and MJT Reinders. Genetic network modeling. Pharmacogenomics, 3(4):507–525, 2002.

[3] Guy Karlebach and Ron Shamir. Modelling and analysis of gene regulatory networks. Nature Reviews Molecular Cell Biology, 9(10):770–780, 2008.

[4] Daniel Marbach, James C Costello, Robert Küffner, Nicole M Vega, Robert J Prill, Diogo M Camacho, Kyle R Allison, Manolis Kellis, James J Collins, Gustavo Stolovitzky, et al. Wisdom of crowds for robust gene network inference. Nature methods, 9(8):796–804, 2012.

[5] Timothy S Gardner and Jeremiah J Faith. Reverse-engineering transcription control networks. Physics of life reviews, 2(1):65–88, 2005.

[6] Jianlong Qi and Tom Michoel. Context-specific transcriptional regulatory network inference from global gene expression maps using double two-way t-tests. Bioinformatics, 28(18):2325–2332, 2012.

[7] Robert J Prill, Daniel Marbach, Julio Saez-Rodriguez, Peter K Sorger, Leonidas G Alexopoulos, Xiaowei Xue, Neil D Clarke, Gregoire Altan-Bonnet, and Gustavo Stolovitzky. Towards a rigorous assessment of systems biology models: the dream3 challenges. PloS one, 5(2):e9202, 2010.

[8] Anne-Claire Haury, Fantine Mordelet, Paola Vera-Licona, and Jean-Philippe Vert. Tigress: trustful inference of gene regulation using stability selection. BMC systems biology, 6(1):1, 2012.

[9] Nir Friedman, Michal Linial, Iftach Nachman, and Dana Pe’er. Using bayesian networks to analyze expression data. Journal of computational biology, 7(3-4):601–620, 2000.

[10] Eran Segal, Haidong Wang, and Daphne Koller. Discovering molecular pathways from protein interaction and gene expression data. Bioinformatics, 19(suppl 1):i264–i272, 2003.

[11] Bruno-Edouard Perrin, Liva Ralaivola, Aurelien Mazurie, Samuele Bottani, Jacques Mallet, and Florence dAlche Buc. Gene networks inference using dynamic bayesian networks. Bioinformatics, 19(suppl 2):ii138–ii148, 2003.

[12] Jing Yu, V Anne Smith, Paul P Wang, Alexander J Hartemink, and Erich D Jarvis. Advances to bayesian network inference for generating causal networks from observational biological data. Bioinformatics, 20(18):3594–3603, 2004.

[13] Michele Ceccarelli, Luigi Cerulo, and Antonella Santone. De novo reconstruction of gene regulatory networks from time series data, an approach based on formal methods. Methods, 69(3):298–305, 2014.

[14] Florian Markowetz and Rainer Spang. Inferring cellular networks–a review. BMC bioinformatics, 8(6):1, 2007.

[15] Jeremiah J Faith, Boris Hayete, Joshua T Thaden, Ilaria Mogno, Jamey Wierzbowski, Guillaume Cottarel, Simon Kasif, James J Collins, and Timothy S Gardner. Large-scale mapping and validation of escherichia coli transcriptional regulation from a compendium of expression profiles. PLoS biol, 5(1):e8, 2007.

[16] Adam A Margolin, Ilya Nemenman, Katia Basso, Chris Wiggins, Gustavo Stolovitzky, Riccardo D Favera, and Andrea Califano. Aracne: an algorithm for the reconstruction of gene regulatory networks in a mammalian cellular context. BMC bioinformatics, 7(Suppl 1):S7, 2006.

[17] Pietro Zoppoli, Sandro Morganella, and Michele Ceccarelli. Timedelay-aracne: Reverse engineering of gene networks from time-course data by an information theoretic approach. BMC bioinformatics, 11(1):154, 2010.

[18] Janusz Sławek and Tomasz Arodź. Ennet: inferring large gene regulatory networks from expression data using gradient boosting. BMC systems biology, 7(1):1, 2013.

[19] Néhémy Lim, Yasin Şenbabaoğlu, George Michailidis, and Florence dAlché Buc. Okvar-boost: a novel boosting algorithm to infer nonlinear dynamics and interactions in gene regulatory networks. Bioinformatics, 29(11):1416–1423, 2013.

[20] Francesca Petralia, Pei Wang, Jialiang Yang, and Zhidong Tu. Integrative random forest for gene regulatory network inference. Bioinformatics, 31(12):i197–i205, 2015.

[21] Alexandre Irrthum, Louis Wehenkel, Pierre Geurts, et al. Inferring regulatory networks from expression data using tree-based methods. PloS one, 5(9):e12776, 2010.

[22] Per Christian Hansen, Toke Koldborg Jensen, and Giuseppe Rodriguez. An adaptive pruning algorithm for the discrete l-curve criterion. Journal of computational and applied mathematics, 198(2):483–492, 2007.

[23] D Calvetti, S Morigi, L Reichel, and F Sgallari. Tikhonov regularization and the l-curve for large discrete ill-posed problems. Journal of computational and applied mathematics, 123(1):423–446, 2000.

[24] L. Garofano, S.M. Pagnotta, and M. Ceccarelli. Synthetic rna-seq network generation and mutual information estimates. https://github.com/lucgar/synRNASeqNet, 2015.

[25] Rob Hyndman and Yeasmin Khandakar. Automatic time series forecasting: The forecast package for r. Journal of Statistical Software, 27(1):1–22, 2008.

[26] Taiwo Adigun, Angela Makolo, and Segun Fatumo. Input dataset survey of in-silico tools for inference and visualization of gene regulatory networks (grn). Computational Biology and Bioinformatics, 3(6):81–87, 2016.

[27] Daniel Marbach, Robert J. Prill, Thomas Schaffter, Claudio Mattiussi, Dario Floreano, and Gustavo Stolovitzky. Revealing strengths and weaknesses of methods for gene network inference. PNAS, 107(14):6286–6291, 2010. WingX.

[28] Anthony Mathelier, Oriol Fornes, David J Arenillas, Chih-yu Chen, Grégoire Denay, Jessica Lee, Wenqiang Shi, Casper Shyr, Ge Tan, Rebecca Worsley-Hunt, et al. Jaspar 2016: a major expansion and update of the open-access database of transcription factor binding profiles. Nucleic acids research, 44(D1):D110–D115, 2016.

[29] Arttu Jolma, Jian Yan, Thomas Whitington, Jarkko Toivonen, Kazuhiro R Nitta, Pasi Rastas, Ekaterina Morgunova, Martin Enge, Mikko Taipale, Gonghong Wei, et al. Dna-binding specificities of human transcription factors. Cell, 152(1):327–339, 2013.

[30] Yue Zhao and Gary D Stormo. Quantitative analysis demonstrates most transcription factors require only simple models of specificity. Nature biotechnology, 29(6):480–483, 2011.

[31] Ivan V Kulakovskiy, Ilya E Vorontsov, Ivan S Yevshin, Anastasiia V Soboleva, Artem S Kasianov, Haitham Ashoor, Wail Ba-Alawi, Vladimir B Bajic, Yulia A Medvedeva, Fedor A Kolpakov, et al. Hocomoco: expansion and enhancement of the collection of transcription factor binding sites models. Nucleic acids research, 44(D1):D116–D125, 2016.

[32] Jason Ernst and Manolis Kellis. Chromhmm: automating chromatin-state discovery and characterization. Nature methods, 9(3):215–216, 2012.

[33] Eugene Tuv, Alexander Borisov, George Runger, and Kari Torkkola. Feature selection with ensembles, artificial variables, and redundancy elimination. Journal of Machine Learning Research, 10(Jul):1341–1366, 2009.

[34] Isabelle Guyon and André Elisseeff. An introduction to variable and feature selection. Journal of machine learning research, 3(Mar):1157–1182, 2003.

[35] Robert Tibshirani. Regression shrinkage and selection via the lasso. Journal of the Royal Statistical Society. Series B (Methodological), pages 267–288, 1996.

[36] Lukas Meier, Sara Van De Geer, and Peter Bühlmann. The group lasso for logistic regression. Journal of the Royal Statistical Society: Series B (Statistical Methodology), 70(1):53–71, 2008.

[37] Robert Tibshirani, Michael Saunders, Saharon Rosset, Ji Zhu, and Keith Knight. Sparsity and smoothness via the fused lasso. Journal of the Royal Statistical Society: Series B (Statistical Methodology), 67(1):91–108, 2005.

[38] Hui Zou and Trevor Hastie. Regularization and variable selection via the elastic net. Journal of the Royal Statistical Society: Series B (Statistical Methodology), 67(2):301–320, 2005.

[39] Nooshin Omranian, Jeanne MO Eloundou-Mbebi, Bernd Mueller-Roeber, and Zoran Nikoloski. Gene regulatory network inference using fused lasso on multiple data sets. Scientific reports, 6, 2016.

[40] Jagath C Rajapakse and Piyushkumar A Mundra. Stability of building gene regulatory networks with sparse autoregressive models. BMC bioinformatics, 12(13):1, 2011.

[41] Andy Liaw and Matthew Wiener. Classification and regression by random-forest. R news, 2(3):18–22, 2002.

[42] Janusz Sławek and Tomasz Arodź. Adanet: inferring gene regulatory networks using ensemble classifiers. In Proceedings of the ACM Conference on Bioinformatics, Computational Biology and Biomedicine, pages 434–441. ACM, 2012.

[43] Per Christian Hansen. The L-curve and its use in the numerical treatment of inverse problems. IMM, Department of Mathematical Modelling, Technical Universityof Denmark, 1999.

[44] Per Christian Hansen and Dianne Prost O’Leary. The use of the l-curve in the regularization of discrete ill-posed problems. SIAM Journal on Scientific Computing, 14(6):1487–1503, 1993.

[45] Per Christian Hansen. Regularization tools: A matlab package for analysis and solution of discrete ill-posed problems. Numerical algorithms, 6(1):1–35, 1994.

[46] Réka Albert and Albert-Lászlό Barabási. Statistical mechanics of complex networks. Reviews of modern physics, 74(1):47, 2002.

[47] J Longina Castellanos, Susana Gόmez, and Valia Guerra. The triangle method for finding the corner of the l-curve. Applied Numerical Mathematics, 43(4):359–373, 2002.

[48] Scott Shaobing Chen and Ramesh A Gopinath. Gaussianization, 2000.

[49] Valero Laparra, Gustavo Camps-Valls, and Jesus Malo. Iterative gaus-sianization: from ica to random rotations. IEEE Transactions on Neural Networks, 22(4):537–549, 2011.

[50] Frank Wilcoxon. Individual comparisons by ranking methods. Biometrics bulletin, 1(6):80–83, 1945.

[51] Robert J Prill, Daniel Marbach, Julio Saez-Rodriguez, Peter K Sorger, Leonidas G Alexopoulos, Xiaowei Xue, Neil D Clarke, Gregoire Altan-Bonnet, and Gustavo Stolovitzky. Towards a rigorous assessment of systems biology models: the dream3 challenges. PloS one, 5(2):e9202, 2010.

[52] Daniel Marbach, Robert J Prill, Thomas Schaffter, Claudio Mattiussi, Dario Floreano, and Gustavo Stolovitzky. Revealing strengths and weaknesses of methods for gene network inference. Proceedings of the national academy of sciences, 107(14):6286–6291, 2010.

[53] Daniel Marbach, Thomas Schaffter, Claudio Mattiussi, and Dario Floreano. Generating realistic in silico gene networks for performance assessment of reverse engineering methods. Journal of computational biology, 16(2):229–239, 2009.

[54] Thomas Schaffter, Daniel Marbach, and Dario Floreano. Genenetweaver: in silico benchmark generation and performance profiling of network inference methods. Bioinformatics, 27(16):2263–2270, 2011.

[55] Kevin Y Yip, Roger P Alexander, Koon-Kiu Yan, and Mark Gerstein. Improved reconstruction of in silico gene regulatory networks by integrating knockout and perturbation data. PloS one, 5(1):e8121, 2010.

[56] Andrea Pinna, Nicola Soranzo, and Alberto De La Fuente. From knockouts to networks: establishing direct cause-effect relationships through graph analysis. PloS one, 5(10):e12912, 2010.

[57] Socorro Gama-Castro, Heladia Salgado, Martin Peralta-Gil, Alberto Santos-Zavaleta, Luis Muniz-Rascado, Hilda Solano-Lira, Veronica Jimenez-Jacinto, Verena Weiss, Jair S Garcia-Sotelo, Alejandra Lopez-Fuentes, et al. Regulondb version 7.0: transcriptional regulation of escherichia coli k-12 integrated within genetic sensory response units (gensor units). Nucleic acids research, 39(suppl 1):D98–D105, 2011.

[58] Michael Ashburner, Catherine A Ball, Judith A Blake, David Botstein, Heather Butler, J Michael Cherry, Allan P Davis, Kara Dolinski, Selina S Dwight, Janan T Eppig, et al. Gene ontology: tool for the unification of biology. Nature genetics, 25(1):25–29, 2000.

[59] Luciano Garofano, Stefano Pagnotta Mario, and Michele Ceccarelli. Synthetic RNA-Seq Network Generation and Mutual Information Estimates. https://github.com/lucgar/synRNASeqNet, 2015.

[60] Norman Lloyd Johnson, Samuel Kotz, and Narayanaswamy Balakrishnan. Discrete multivariate distributions, volume 165. Wiley New York, 1997.

[61] Michal Ronen, Revital Rosenberg, Boris I Shraiman, and Uri Alon. Assigning numbers to the arrows: parameterizing a gene regulation network by using accurate expression kinetics. Proceedings of the national academy of sciences, 99(16):10555–10560, 2002.

[62] Paul T Spellman, Gavin Sherlock, Michael Q Zhang, Vishwanath R Iyer, Kirk Anders, Michael B Eisen, Patrick O Brown, David Botstein, and Bruce Futcher. Comprehensive identification of cell cycle-regulated genes of the yeast saccharomyces cerevisiae by microarray hybridization. Molecular biology of the cell, 9(12):3273–3297, 1998.

[63] Irene Cantone, Lucia Marucci, Francesco Iorio, Maria Aurelia Ricci, Vincenzo Belcastro, Mukesh Bansal, Stefania Santini, Mario Di Bernardo, Diego Di Bernardo, and Maria Pia Cosma. A yeast synthetic network for in vivo assessment of reverse-engineering and modeling approaches. Cell, 137(1):172–181, 2009.

[64] Michele Ceccarelli, Luigi Cerulo, and Antonella Santone. De novo reconstruction of gene regulatory networks from time series data, an approach based on formal methods. Methods, 69(3):298–305, 2014.

[65] Athanasios Polynikis, SJ Hogan, and Mario di Bernardo. Comparing different ode modelling approaches for gene regulatory networks. Journal of theoretical biology, 261(4):511–530, 2009.

[66] P. Y. Wen and S. Kesari. Malignant gliomas in adults. N. Engl. J. Med., 359(5):492–507, Jul 2008.

[67] Maria Stella Carro, Wei Keat Lim, Mariano Javier Alvarez, Robert J Bollo, Xudong Zhao, Evan Y Snyder, Erik P Sulman, Sandrine L Anne, Fiõna Doetsch, Howard Colman, et al. The transcriptional network for mesenchymal transformation of brain tumours. Nature, 463(7279):318–325, 2010.

[68] Celine Lefebvre, Presha Rajbhandari, Mariano J Alvarez, Pradeep Bandaru, Wei Keat Lim, Mai Sato, Kai Wang, Pavel Sumazin, Manjunath Kustagi, Brygida C Bisikirska, et al. A human b-cell interactome identifies myb and foxm1 as master regulators of proliferation in germinal centers. Molecular systems biology, 6(1):377, 2010.

[69] Michele Ceccarelli, Floris P Barthel, Tathiane M Malta, Thais S Sabedot, Sofie R Salama, Bradley A Murray, Olena Morozova, Yulia Newton, Amie Radenbaugh, Stefano M Pagnotta, et al. Molecular profiling reveals biologically discrete subsets and pathways of progression in diffuse glioma. Cell, 164(3):550–563, 2016.

[70] Houtan Noushmehr, Daniel J Weisenberger, Kristin Diefes, Heidi S Phillips, Kanan Pujara, Benjamin P Berman, Fei Pan, Christopher E Pelloski, Erik P Sulman, Krishna P Bhat, et al. Identification of a cpg island methylator phenotype that defines a distinct subgroup of glioma. Cancer cell, 17(5):510–522, 2010.

[71] Raghvendra Mall, Luigi Cerulo, Halima Bensmail, Antonio Iavarone, and Michele Ceccarelli. Detection of statistically significant network changes in complex biological networks. BMC Systems Biology, 11(1):32, 2017.

[72] W Evan Johnson, Cheng Li, and Ariel Rabinovic. Adjusting batch effects in microarray expression data using empirical bayes methods. Biostatistics, 8(1):118–127, 2007.

